## Supplementary Material for "Identifying images in the biology literature that are problematic for people with a color-vision deficiency"

### Supplementary Tables

**Table S1: Predictive performance for metrics that characterize potentially problematic aspects of images.** We calculated five metrics, as well as a rank-based, combined score and assessed their ability to categorize images as “Definitely okay” or “Definitely problematic”.

| **Metric** | **AUROC** | **AUPRC** |
| --- | --- | --- |
| Mean, pixel-wise color distance between the original and simulated image | 0.44 | 0.12 |
| Color-distance ratio between the original and simulated images for the color pair with the largest distance in the original image | 0.63 | 0.19 |
| Number of color pairs that exhibited a high color-distance ratio between the original and simulated images | 0.75 | 0.34 |
| Proportion of pixels in the original image that used a color from one of the high-ratio color pairs | 0.73 | 0.28 |
| Mean Euclidean distance between pixels for high-ratio color pairs | 0.67 | 0.24 |
| Rank-based score that combines the metrics | 0.71 | 0.26 |

**Table S2: Predictive performance for classification algorithms that used the image metrics as inputs.** We used classification algorithms to categorize images as “Definitely okay” or “Definitely problematic”. These results indicate the algorithms’ performance after cross validation on the training set.

| **Algorithm** | **AUROC** | **AUPRC** |
| --- | --- | --- |
| Logistic Regression | 0.82 | 0.43 |
| Nearest Neighbors | 0.72 | 0.32 |
| Random Forests | 0.80 | 0.42 |

**Table S3: Predictive performance for Convolutional Neural Network models that used the images as inputs.** We tested 23 model configurations via cross validation on the training set, evaluating each model’s ability to categorize images as “Definitely okay” or “Definitely problematic”.

| **Combination** | **Class weighting** | **Early stopping** | **Random rotation** | **Dropout** | **Transfer learning** | **Fine tuning** | **AUROC** | **AUPRC** |
| --- | --- | --- | --- | --- | --- | --- | --- | --- |
| 0 | No | No | 0.0 | 0.0 | None | No | 0.77 | 0.48 |
| 1 | Yes | No | 0.0 | 0.0 | None | No | 0.83 | 0.51 |
| 2 | No | Yes | 0.0 | 0.0 | None | No | 0.88 | 0.58 |
| 3 | No | No | 0.2 | 0.0 | None | No | 0.91 | 0.68 |
| 4 | No | No | 0.3 | 0.0 | None | No | 0.91 | 0.68 |
| 5 | No | No | 0.0 | 0.2 | None | No | 0.78 | 0.51 |
| 6 | No | No | 0.0 | 0.5 | None | No | 0.80 | 0.45 |
| 7 | No | No | 0.0 | 0.0 | MobileNetV2 | No | 0.85 | 0.55 |
| 8 | No | No | 0.0 | 0.0 | ResNet50 | No | 0.87 | 0.62 |
| 9 | No | No | 0.0 | 0.0 | MobileNetV2 | Yes | 0.85 | 0.61 |
| 10 | No | No | 0.0 | 0.0 | ResNet50 | Yes | 0.85 | 0.62 |
| 11 | Yes | Yes | 0.2 | 0.0 | ResNet50 | No | 0.89 | 0.62 |
| 12 | Yes | Yes | 0.2 | 0.2 | ResNet50 | No | 0.88 | 0.60 |
| 13 | Yes | Yes | 0.2 | 0.5 | ResNet50 | No | 0.88 | 0.58 |
| 14 | Yes | Yes | 0.2 | 0.0 | ResNet50 | Yes | 0.92 | 0.74 |
| 15 | Yes | Yes | 0.2 | 0.2 | ResNet50 | Yes | 0.92 | 0.74 |
| 16 | Yes | Yes | 0.2 | 0.5 | ResNet50 | Yes | 0.93 | 0.75 |
| 17 | Yes | Yes | 0.2 | 0.0 | MobileNetV2 | No | 0.86 | 0.52 |
| 18 | Yes | Yes | 0.2 | 0.2 | MobileNetV2 | No | 0.86 | 0.52 |
| 19 | Yes | Yes | 0.2 | 0.5 | MobileNetV2 | No | 0.87 | 0.53 |
| 20 | Yes | Yes | 0.2 | 0.0 | MobileNetV2 | Yes | 0.89 | 0.65 |
| 21 | Yes | Yes | 0.2 | 0.2 | MobileNetV2 | Yes | 0.90 | 0.68 |
| 22 | Yes | Yes | 0.2 | 0.5 | MobileNetV2 | Yes | 0.91 | 0.69 |

### Supplementary Figures

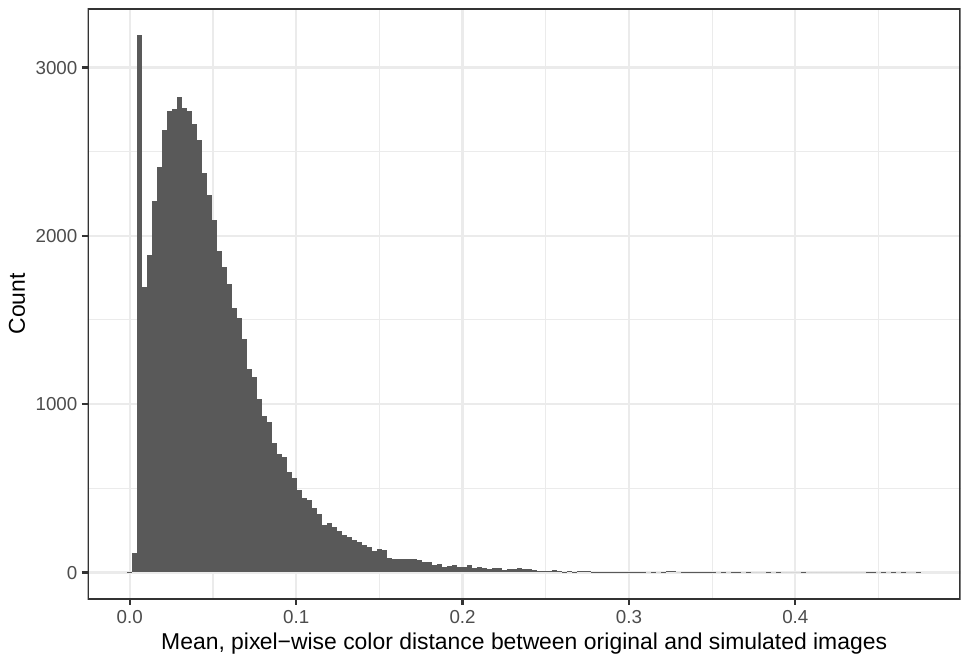

**Figure S1: Mean, pixel-wise color distance between each original and simulated image from *eLife*.** The histogram depicts the frequency distribution of this metric for 64,509 non-grayscale images.

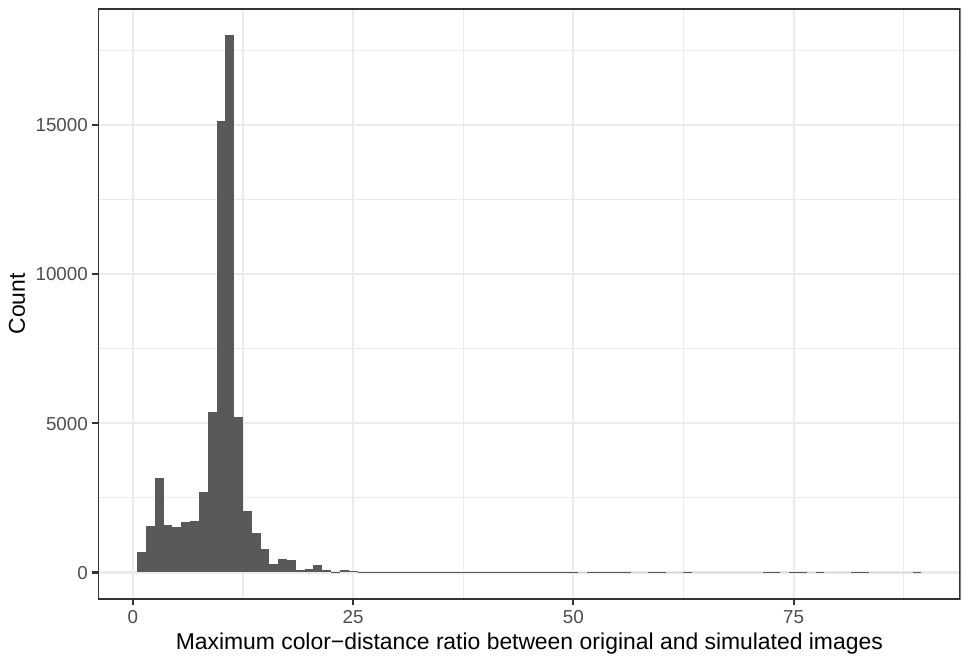

**Figure S2: Maximum color-distance ratio between each original and simulated image from *eLife*.** The histogram depicts the frequency distribution of this metric for 64,509 non-grayscale images.

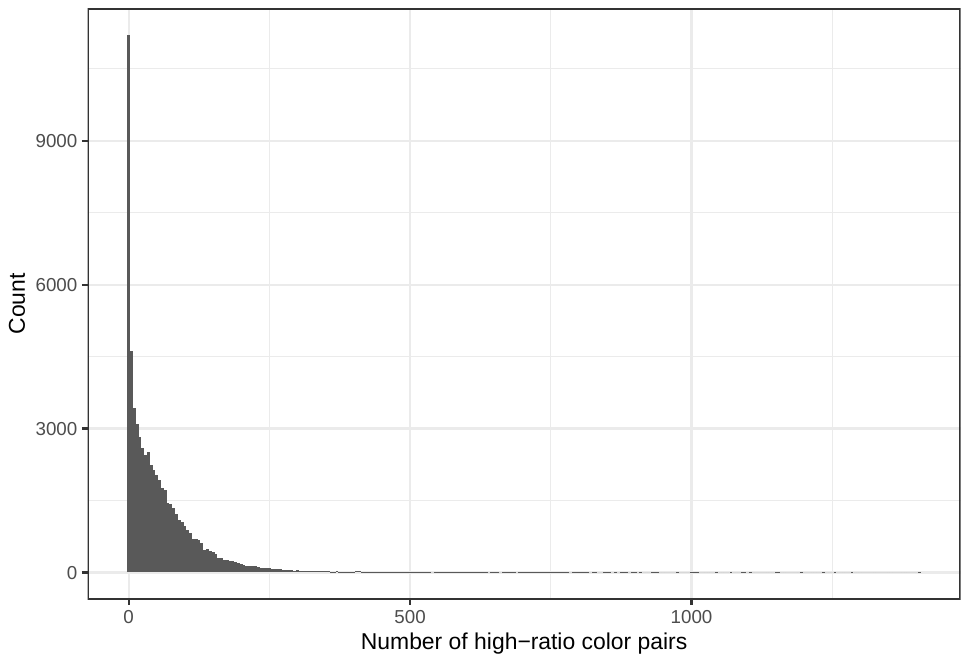

**Figure S3: Number of color pairs per image that exhibited a high color-distance ratio between the original and simulated images from *eLife*.** The histogram depicts the frequency distribution of this metric for 64,509 non-grayscale images.

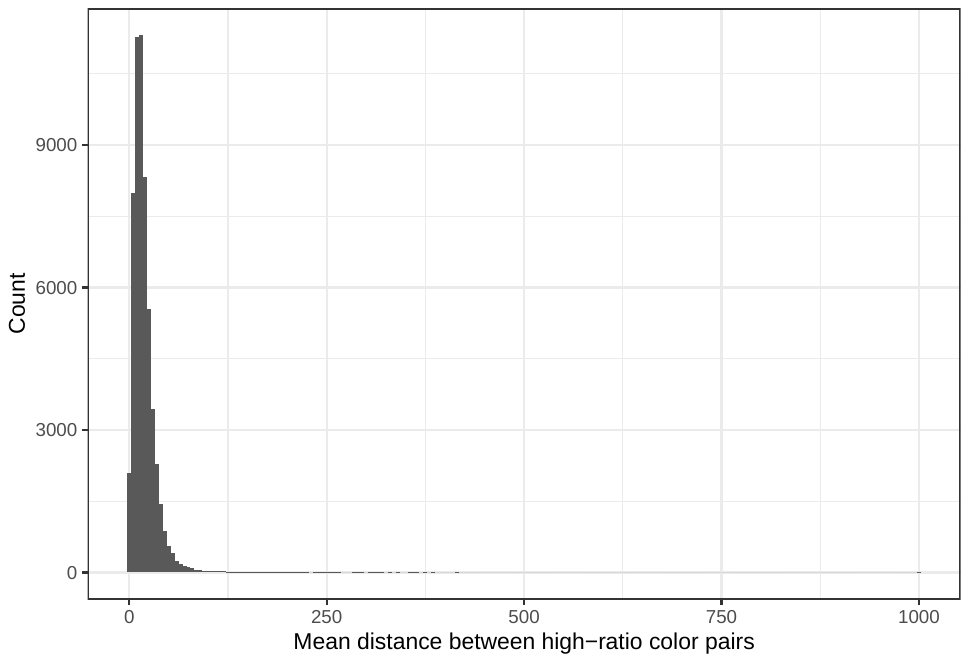

**Figure S4: Mean Euclidean spatial distance per image between pixels for high-ratio color pairs from *eLife*.** The histogram depicts the frequency distribution of this metric for 64,509 non-grayscale images.

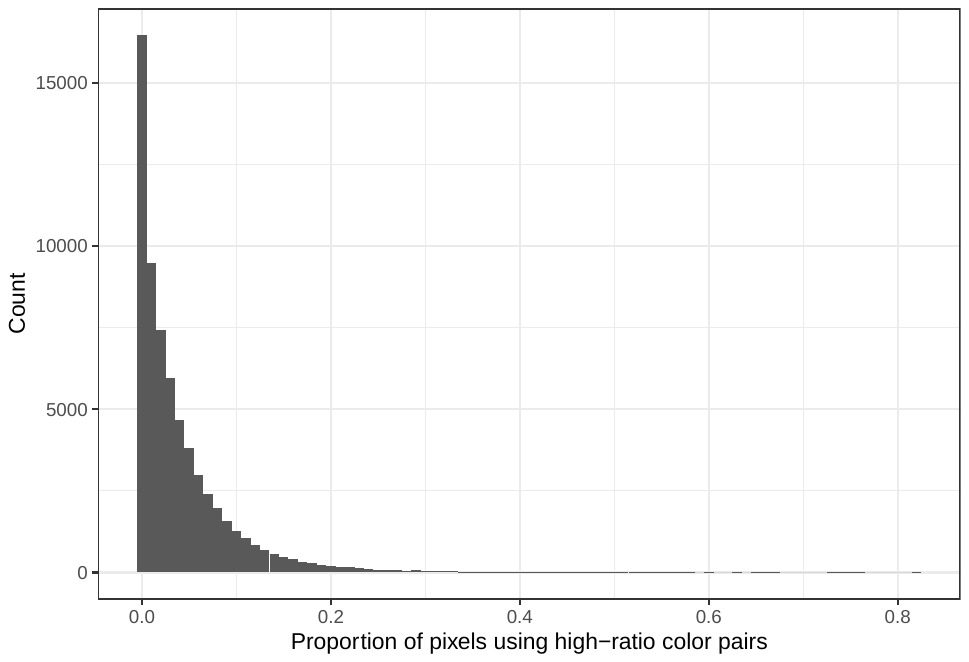

**Figure S5: Proportion of pixels in each original image that used a color from one of the high-ratio color pairs from *eLife*.** The histogram depicts the frequency distribution of this metric for 64,509 non-grayscale images.

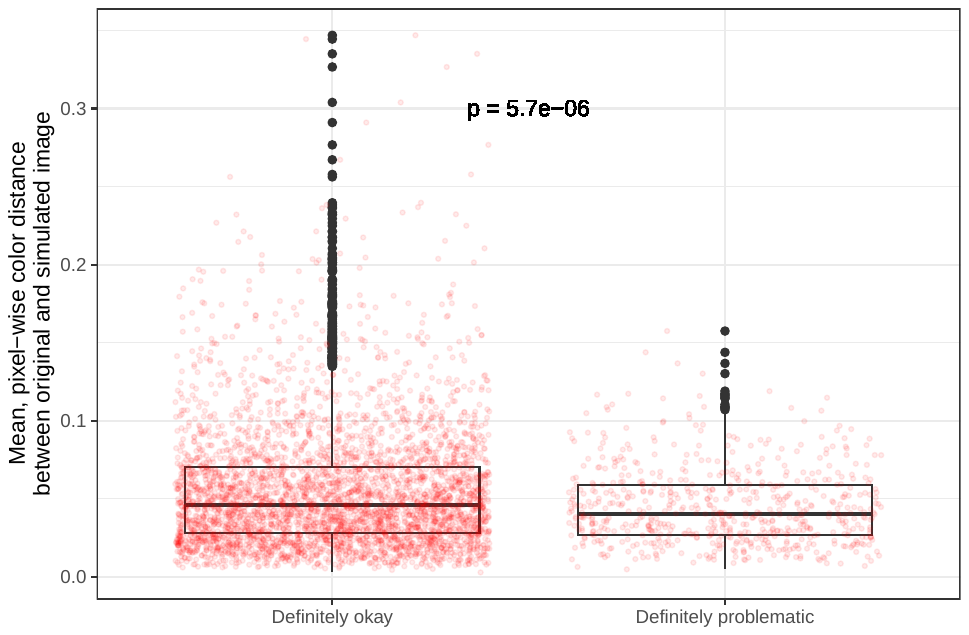

**Figure S6: Mean, pixel-wise color distance between each original and simulated image from *eLife* categorized as “Definitely okay” or “Definitely problematic”.** We compared a two-sided Mann-Whitney U test to calculate the p-value.

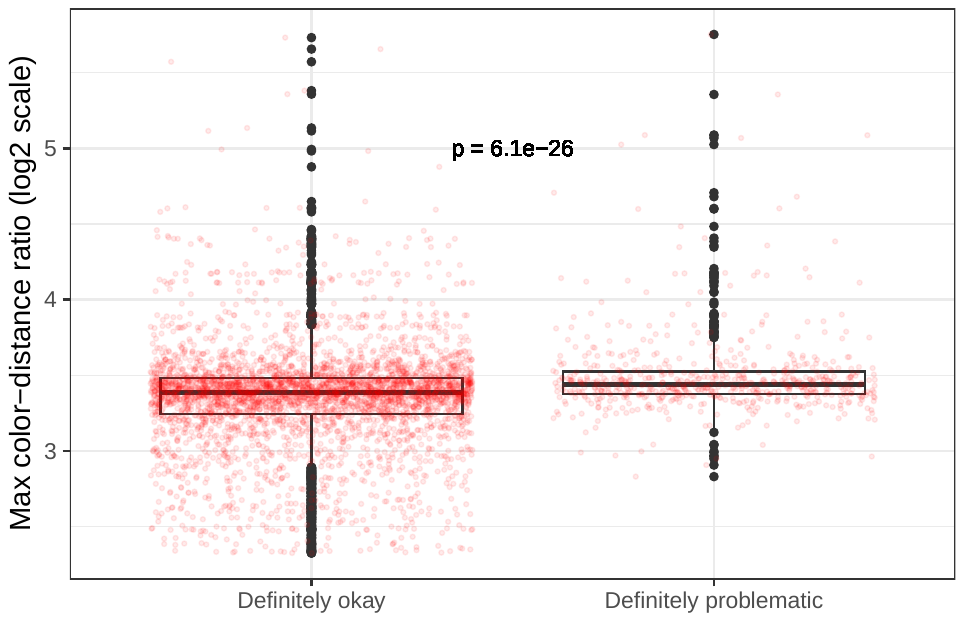

**Figure S7: Maximum color-distance ratio between each original and simulated image from *eLife* categorized as “Definitely okay” or “Definitely problematic”.** We compared a two-sided Mann-Whitney U test to calculate the p-value.

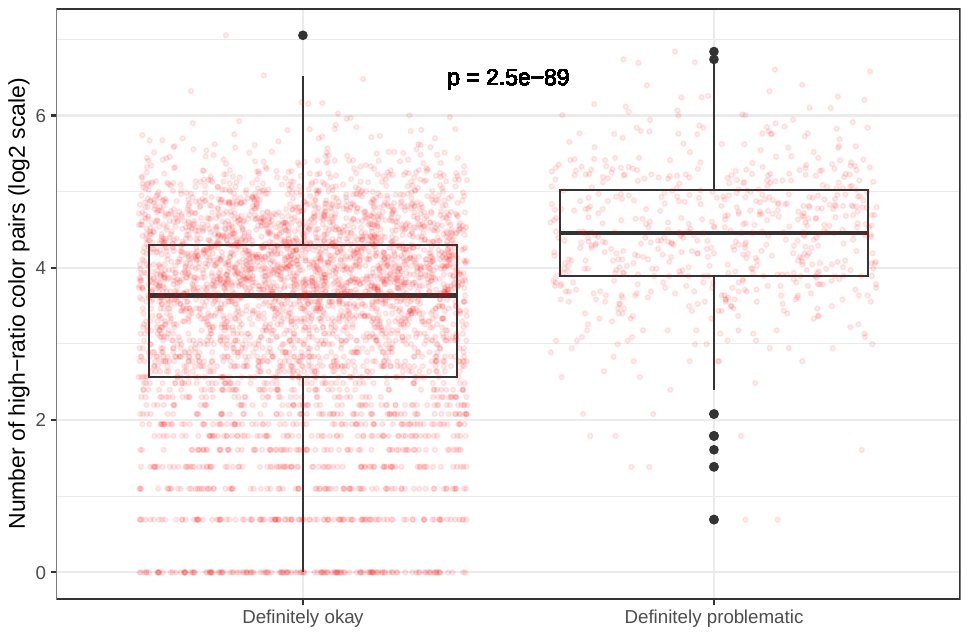

**Figure S8: Number of high-ratio color pairs per image from *eLife* categorized as “Definitely okay” or “Definitely problematic”.** We compared a two-sided Mann-Whitney U test to calculate the p-value.

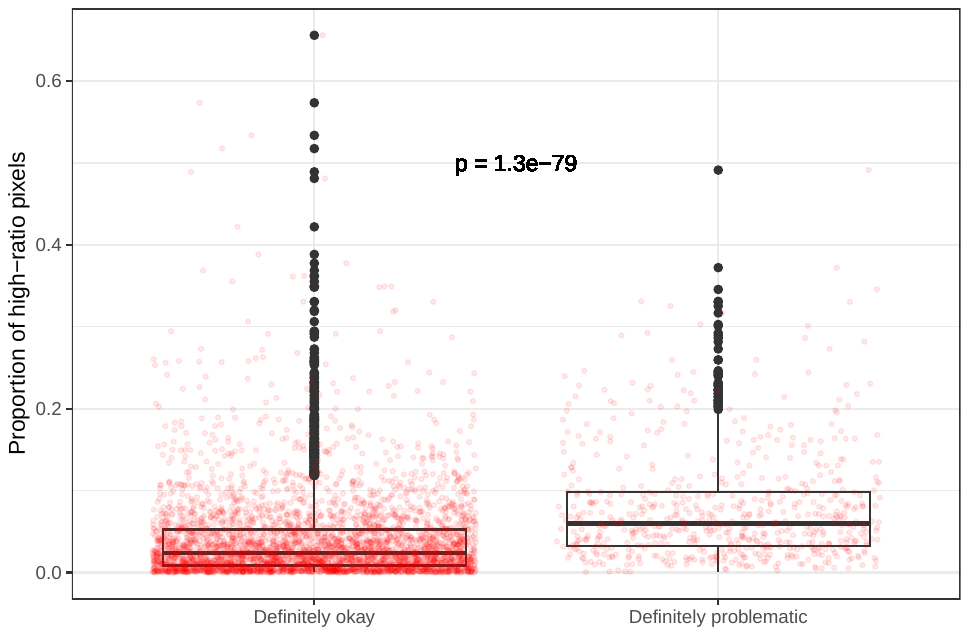

**Figure S9: Proportion of pixels for high-ratio color pairs for images from *eLife* categorized as “Definitely okay” or “Definitely problematic”.** We compared a two-sided Mann-Whitney U test to calculate the p-value.

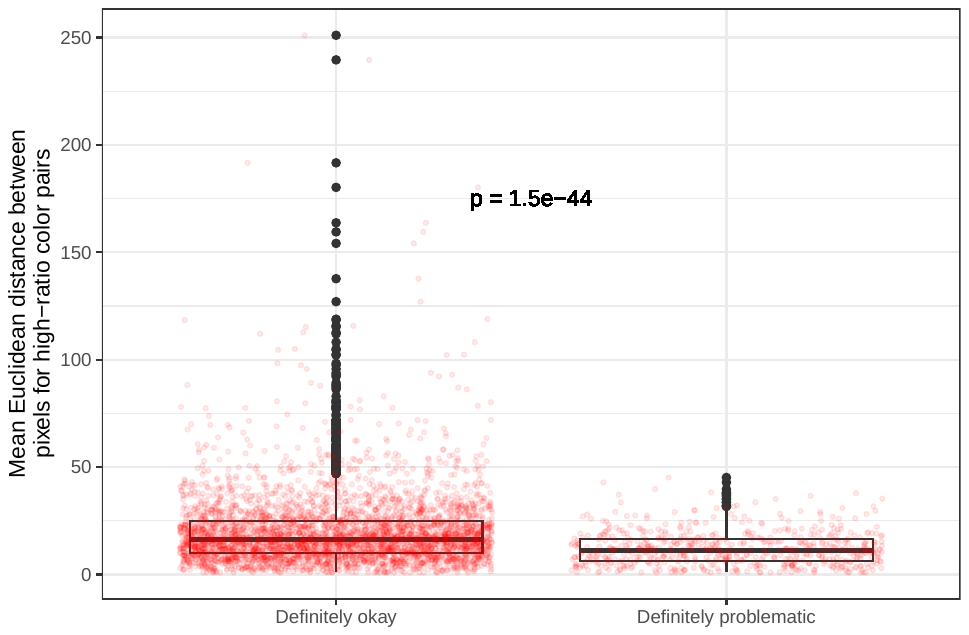

**Figure S10: Mean, pixel-wise Euclidean distance for high-ratio color pairs in images from *eLife* categorized as “Definitely okay” or “Definitely problematic”.** We compared a two-sided Mann-Whitney U test to calculate the p-value.

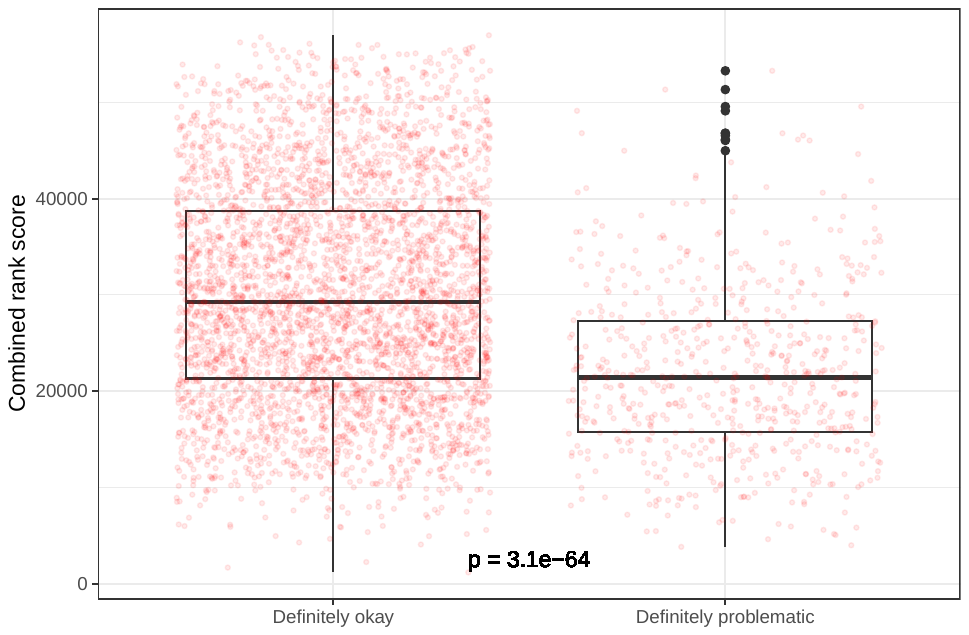

**Figure S11: Rank-based metric score for images from *eLife* categorized as “Definitely okay” or “Definitely problematic”.** We compared a two-sided Mann-Whitney U test to calculate the p-value.

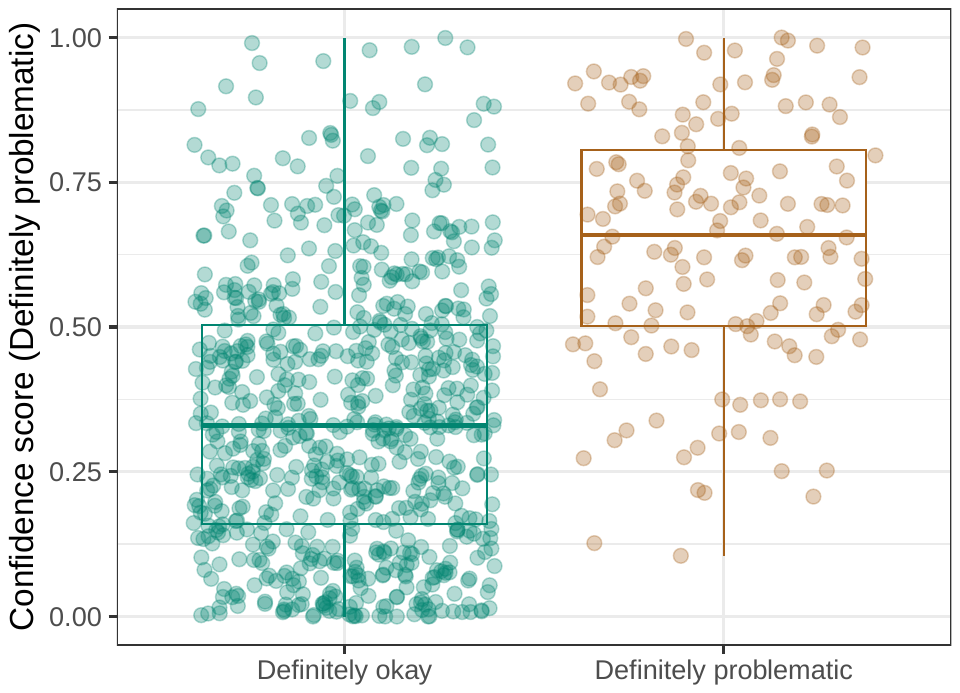

**Figure S12: Logistic Regression predictions for the images in the *eLife* hold-out test set.** Each point represents the prediction for an individual image. Relatively high confidence scores indicate that the model had more confidence that a given image was “Definitely problematic” for a person with deuteranopia.

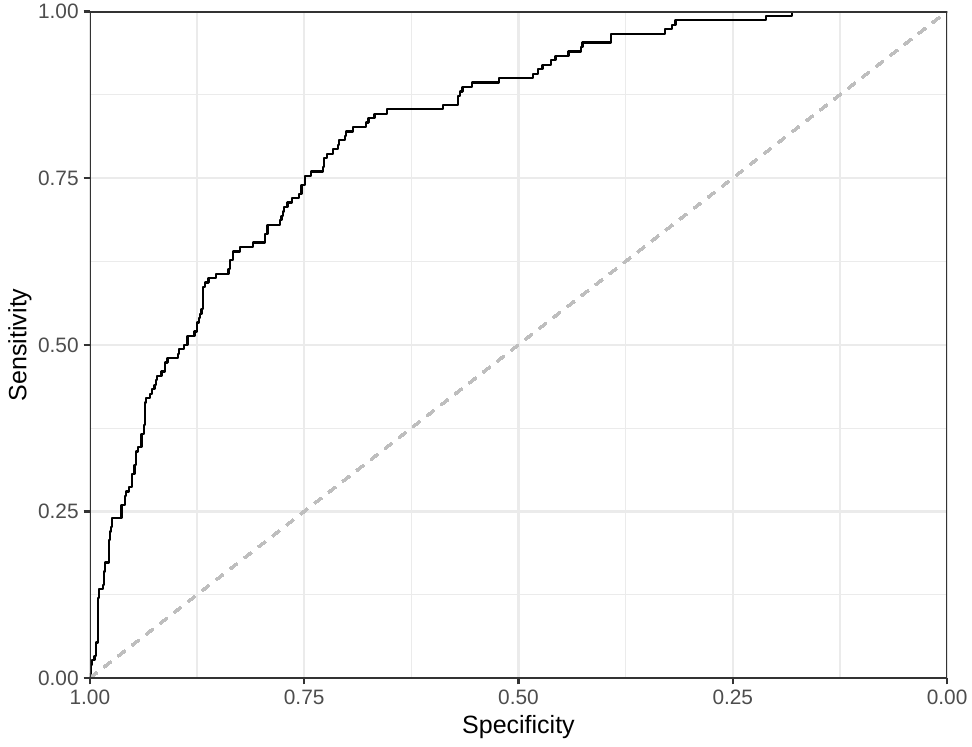

**Figure S13: Receiver operating characteristic curve for the Logistic Regression predictions for the images in the *eLife* hold-out test set.** This curve illustrates tradeoffs between sensitivity and specificity. The area under the curve is 0.82.

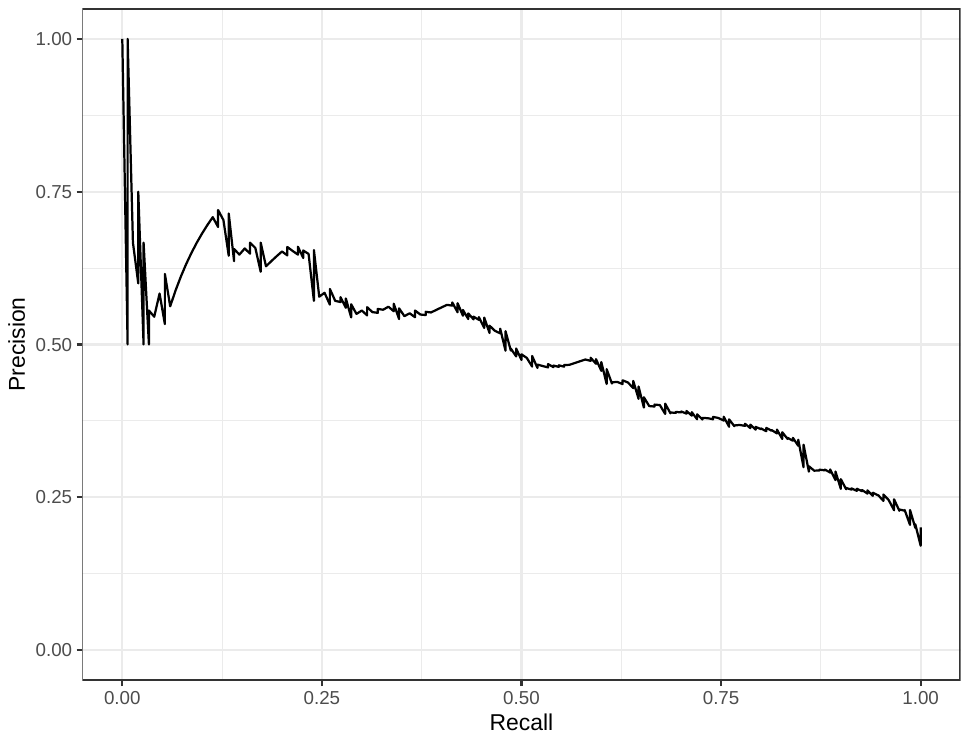

**Figure S14: Precision-recall curve for the Logistic Regression predictions for the images in the *eLife* hold-out test set.** This curve illustrates tradeoffs between precision and recall. The area under the curve is 0.49.

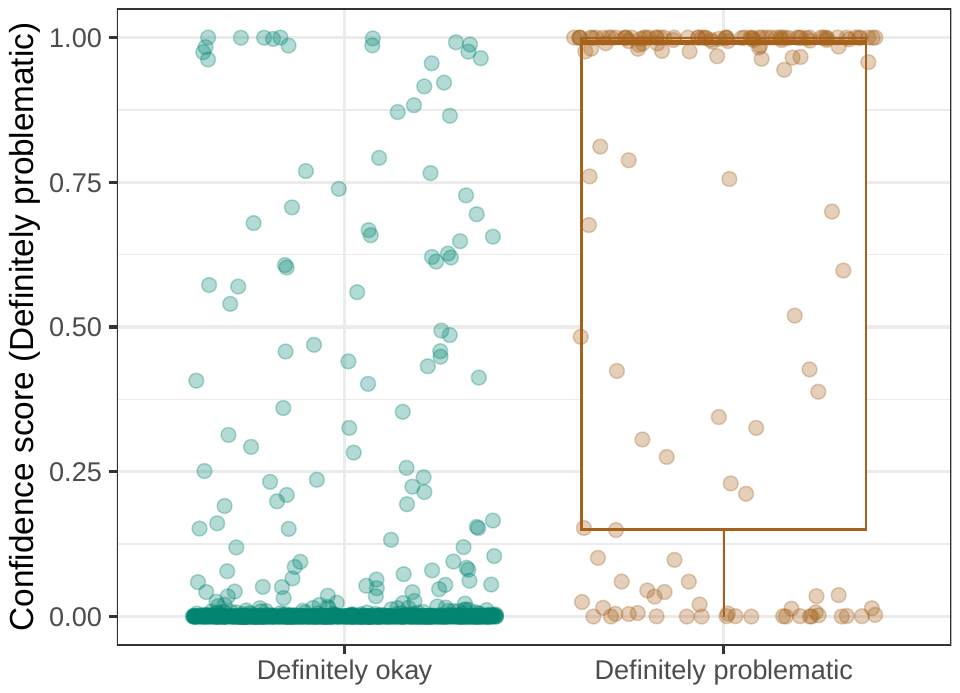

**Figure S15: Convolutional Neural Network predictions for images in the *eLife* hold-out test set.** Each point represents the prediction for an individual image. Relatively high confidence scores indicate that the model had more confidence that a given image was “Definitely problematic” for a person with deuteranopia.

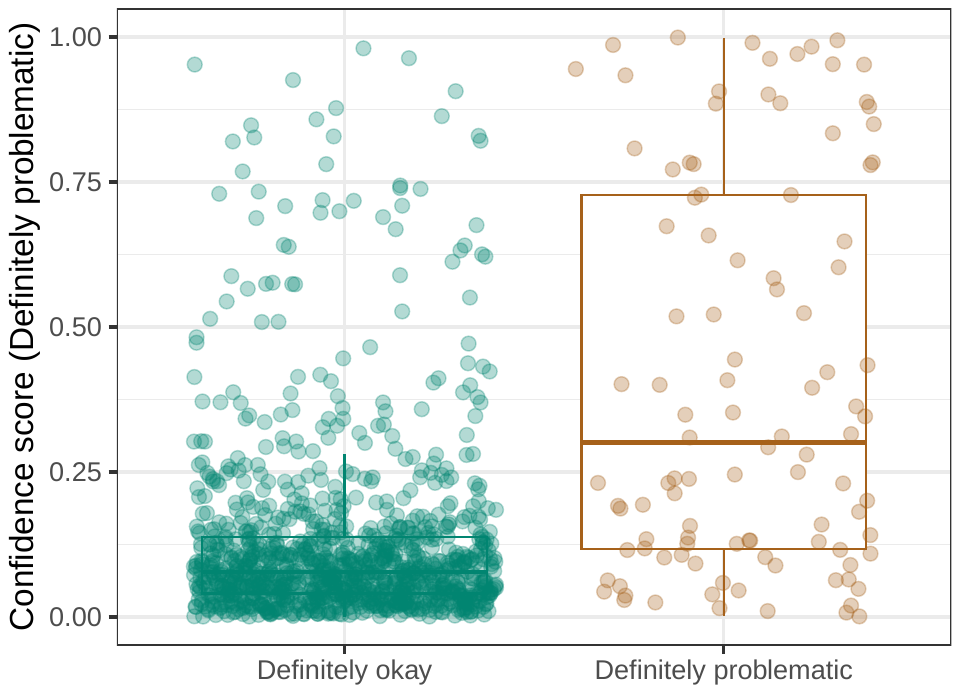

**Figure S16: Convolutional Neural Network predictions for images in the PubMed Central hold-out test set.** Each point represents the prediction for an individual image. Relatively high confidence scores indicate that the model had more confidence that a given image was “Definitely problematic” for a person with deuteranopia.

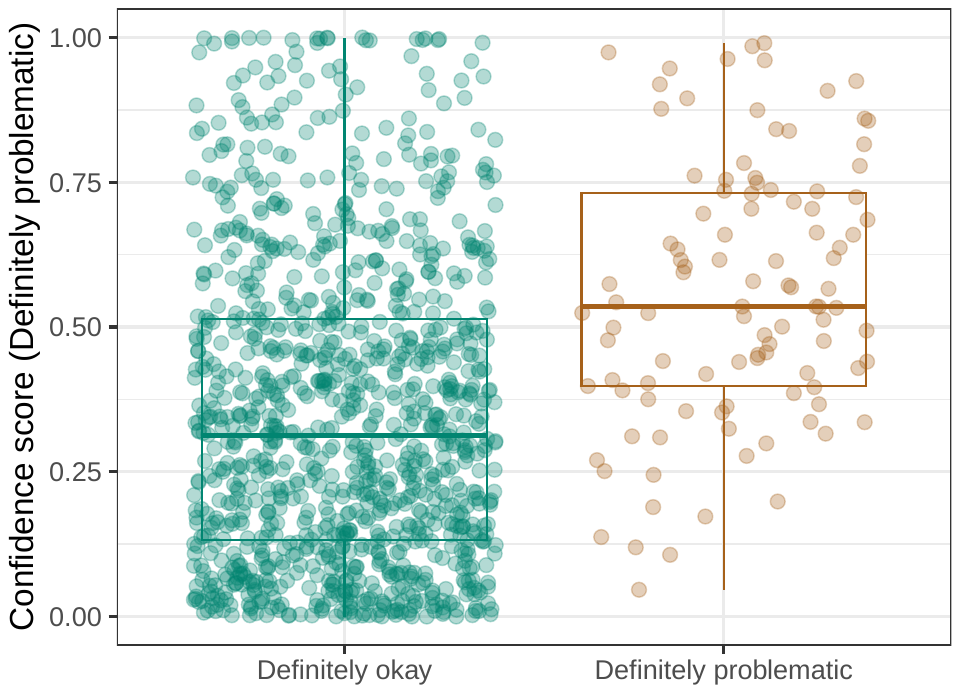

**Figure S17: Logistic Regression predictions for the images in the PubMed Central hold-out test set.** Each point represents the prediction for an individual image. Relatively high confidence scores indicate that the model had more confidence that a given image was “Definitely problematic” for a person with deuteranopia.

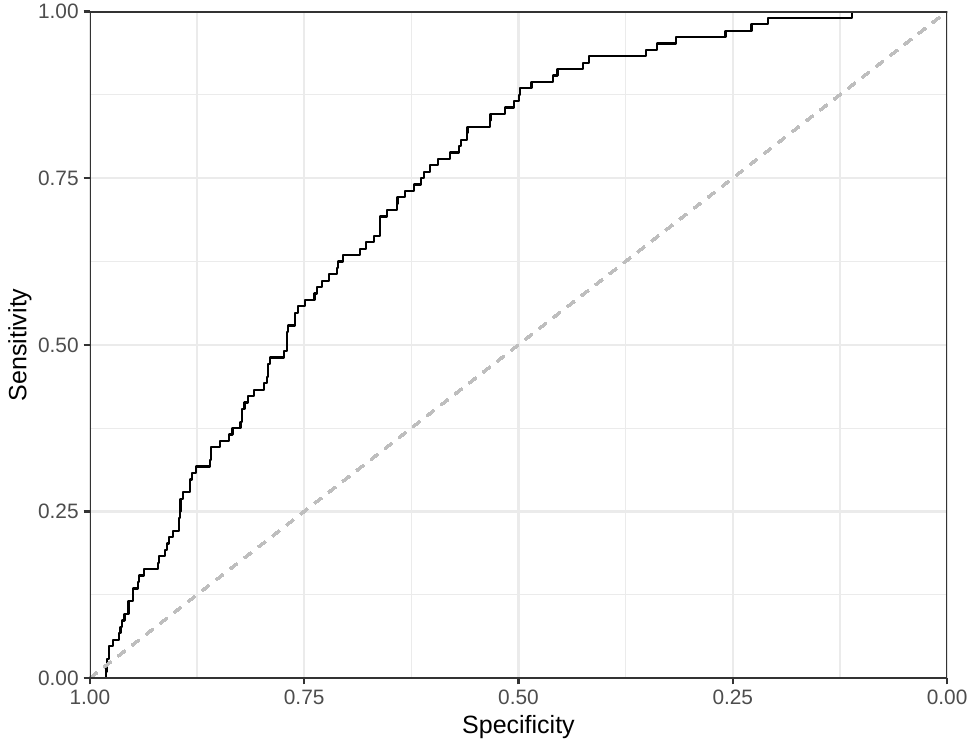

**Figure S18: Receiver operating characteristic curve for the Logistic Regression predictions for the images in the PubMed Central hold-out test set.** This curve illustrates tradeoffs between sensitivity and specificity. The area under the curve is 0.73.

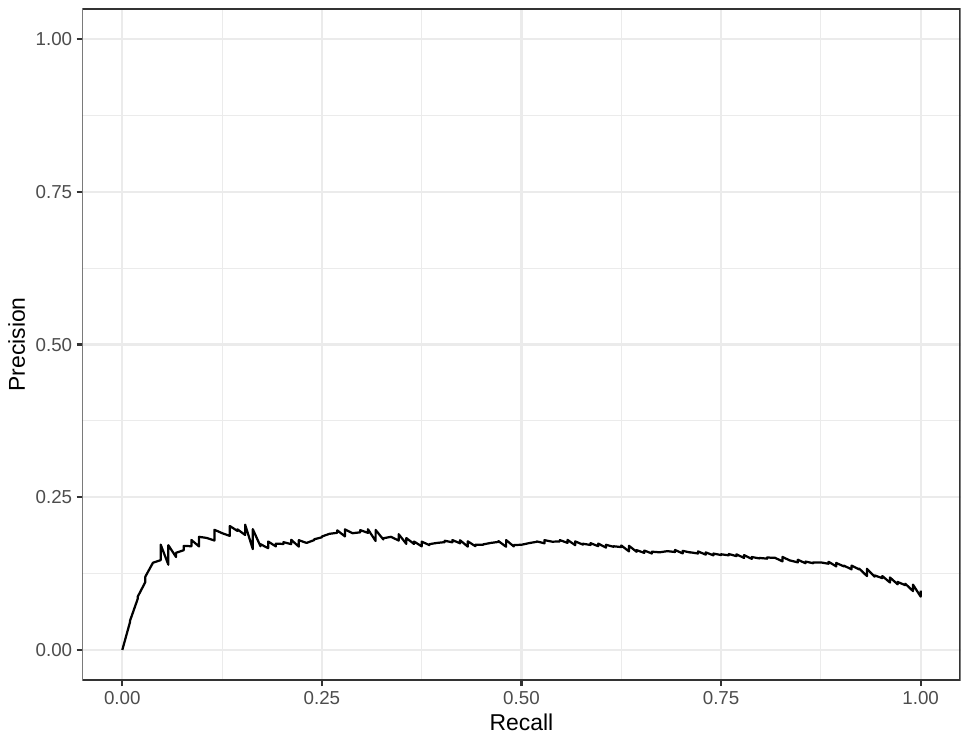

**Figure S19: Precision-recall curve for the Logistic Regression predictions for the images in the PubMed Central hold-out test set.** This curve illustrates tradeoffs between precision and recall. The area under the curve is 0.16.
